# Dynamics of brainstem arousal systems and pupil size predict cortical interactions for flexible decision-making

**DOI:** 10.1101/2023.12.05.570327

**Authors:** R.L. van den Brink, K. Hagena, N. Wilming, P.R. Murphy, J. Calder-Travis, J. Finsterbusch, C. Büchel, T.H. Donner

## Abstract

Most perceptual decisions entail a flexible mapping from sensory input to motor output. Flexible input-output mapping is reflected in correlated variability within the cortical network involved in perceptual decisions. Here, we tested the idea that brainstem arousal systems are involved in flexible input-output mapping and changes in cortical correlated variability. We combined brainstem fMRI, pupillometry, and time-resolved assessment of the intrinsic correlations between cortical population codes for stimulus and action. Human participants reported the orientation of visual stimuli by button presses, while the required stimulus-response mapping rule could undergo hidden and unpredictable changes. Rule switches evoked brainstem and pupil responses as well as changes in computational model-inferred, latent variables. These variables governed participants’ rule-switching behavior and pupil responses. Brainstem activity and pupil dilation preceded increases in the strength of correlations between cortical stimulus and action codes. Brainstem arousal systems may promote the reorganization of sensorimotor cortical pathways for flexible decisions.

## Introduction

Decision-making involves the transformation of sensory signals into motor actions (Mante et al., 2013; Shadlen and Kiani, 2013). The transformation from sensation to action has been studied predominantly using tasks with static associations between stimulus and action (Bogacz et al., 2006; Gold and Shadlen, 2007; Donner et al., 2009; Hanks et al., 2015; Wilming et al., 2020; Murphy et al., 2021). However, the mapping between stimulus and action is often not fixed, but must be flexibly altered in accordance with environmental demands (Okazawa and Kiani, 2023). For example, in order to alternate between two stimulus-response (SR) mapping rules (Figure 1a), the brain must flexibly route stimulus information from visual cortex to the population of neurons in motor cortex that encode the correct action for a given stimulus and SR rule (Miller and Cohen, 2001) (Figure 1b). How the brain is able to flexibly remap SR-associations has remained a key question in decision neuroscience (Shadlen and Kiani, 2013).

**Figure 1.**
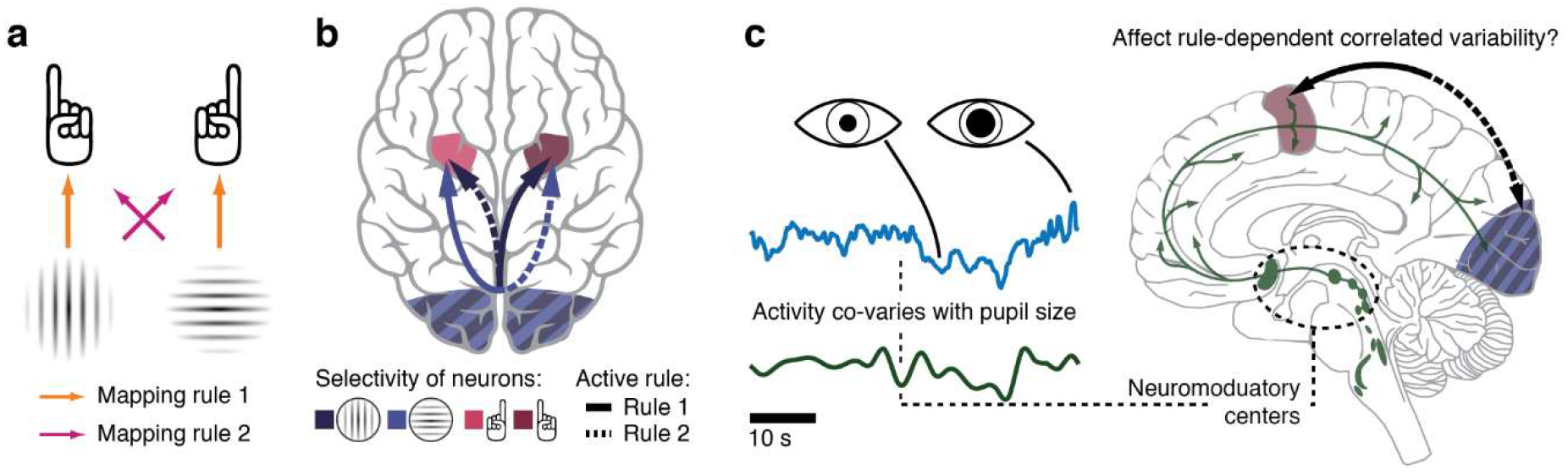
Rationale. **a**. Elementary perceptual choice task. Each visual grating is associated with a unique correct action, which depends on a currently active response rule. **b**. Schematic of rule-dependent correlated variability. Arrows indicate positive correlation. When rule 1 is active, the activity of the population of neurons in visual cortex that encodes a vertical grating correlates positively with those neurons that encode a left hand response. Similarly, horizontal-encoding neurons correlate with the right hand response. When rule 2 becomes active, this pattern flips sign. c. Experimental hypothesis. We expect signals from neuromodulatory brainstem nuclei (and their correlate in pupil responses) to (i) track changes in SR rule and (ii) modulate the strength of features-specific correlated variability in the sensory-motor network.

We recently showed that arbitrary SR rules (Figure 1a) are instantiated in correlated variability of ongoing fluctuations of stimulus and action patterns amongst the regions that implement the decision process (van den Brink et al., 2023). That is, spontaneous fluctuations of orientation-selective fMRI signals in visual cortex were correlated with fluctuations in motor cortex that were selective to the appropriate action (in line with the schematic in Figure 1b). This held both in situations where the SR rule was explicitly instructed and where it switched in a hidden and unpredictable manner and needed to be inferred from noisy cues. In the latter case, the SR coupling patterns reflected the participants’ internal belief about the active SR rule (gauged through a computational model). This observation supports the idea that task-specific SR pathways are constantly reconfigured when dictated by the environment, guided by higher-order inference processes. What mechanisms confer this flexible reconfiguration of SR pathways?

Theoretical work indicates that the brain can flexibility reconfigure SR pathways through short term synaptic plasticity mechanisms that are in turn governed by subcortical neuromodulatory inputs (Fusi et al., 2007). These nuclei project widely to the forebrain and are thus ideally situated to affect large-scale cortical activity dynamics (Berridge and Waterhouse, 2003; Aston-Jones and Cohen, 2005; van den Brink et al., 2019; Pfeffer et al., 2021; Podvalny et al., 2021; Pfeffer et al., 2022), and control cortical state and arousal (Harris and Thiele, 2011; McGinley et al., 2015b). Moreover, neuromodulators are known to promote synaptic plasticity on multiple temporal scales (Bear and Singer, 1986; Rasmusson, 2000; Reynolds et al., 2001; Reynolds and Wickens, 2002; Berridge and Waterhouse, 2003; Vetencourt et al., 2008; Marzo et al., 2009; Nadim and Bucher, 2014). The activity of neurons that release neuromodulators is also sensitive to stimulus features that can signal the need to adjust behavior (Berridge and Waterhouse, 2003; Aston-Jones and Cohen, 2005; Bouret and Sara, 2005; Sarter et al., 2009), including specific computational variables entailed in complex inference strategies (Dayan and Yu, 2006; Nassar et al., 2012; Muller et al., 2019; Murphy et al., 2021). Combined, these characteristics indicate that neuromodulators may track hidden changes in SR rules and sculpt correlated variability of cortical signals that conditionally link stimulus with action (Figure 3c).

We tested these hypotheses by reanalyzing fMRI data from a previously published study (van den Brink et al., 2023). Here, we probe the relationship between the SR rule inference process, and brainstem fMRI signals, pupil diameter, and intrinsic correlations within the human sensory-motor network. Participants carried out an elementary perceptual choice task (Figure 1a) that was coupled to an inference problem: an active SR rule had to be inferred from ambiguous sensory evidence, and underwent hidden and unpredictable switches (van den Brink et al., 2023). The hidden rule switches evoked robust responses in several neuromodulatory nuclei as well as in the pupil (Figure 1c). These physiological markers of arousal in response to rule switches could be understood in terms of a concomitant increase in computational model variables, which signaled that a change in rule was likely to have occurred. These same markers of arousal also preceded an increase in SR coupling strength within the cortical network that implemented the decision. Our results suggest that brainstem arousal is driven by key computational variables for context inference, and helps shape the continuous and context-dependent reorganization of task-specific pathways within the cortex.

## Results

### Inferring volatile sensory-motor mapping rules under uncertainty

Participants (N=18) performed a visual orientation discrimination task in which the stimulus-response mapping rule switched in a hidden and unpredictable fashion. On each trial of the primary task, a vertical or horizontal visual grating (full contrast; see Methods) was presented. Participants reported the grating orientation with a button press, using either the left or right hand. Two distinct rules could govern the required mapping from stimulus orientations to motor effectors, and participants needed to infer the rule that was active at any time from a rapid stream of noisy sensory evidence (Figure 2a). The evidence samples were small dots presented at varying locations on the horizontal meridian, drawn from one of two distributions with different means. The distribution generating the samples was coupled to the active rule, and it could switch unpredictably from sample to sample with a low probability (p = 1/70). The trials of the orientation judgement were interspersed in the stream of evidence samples at long and variable intervals.

**Figure 2.**
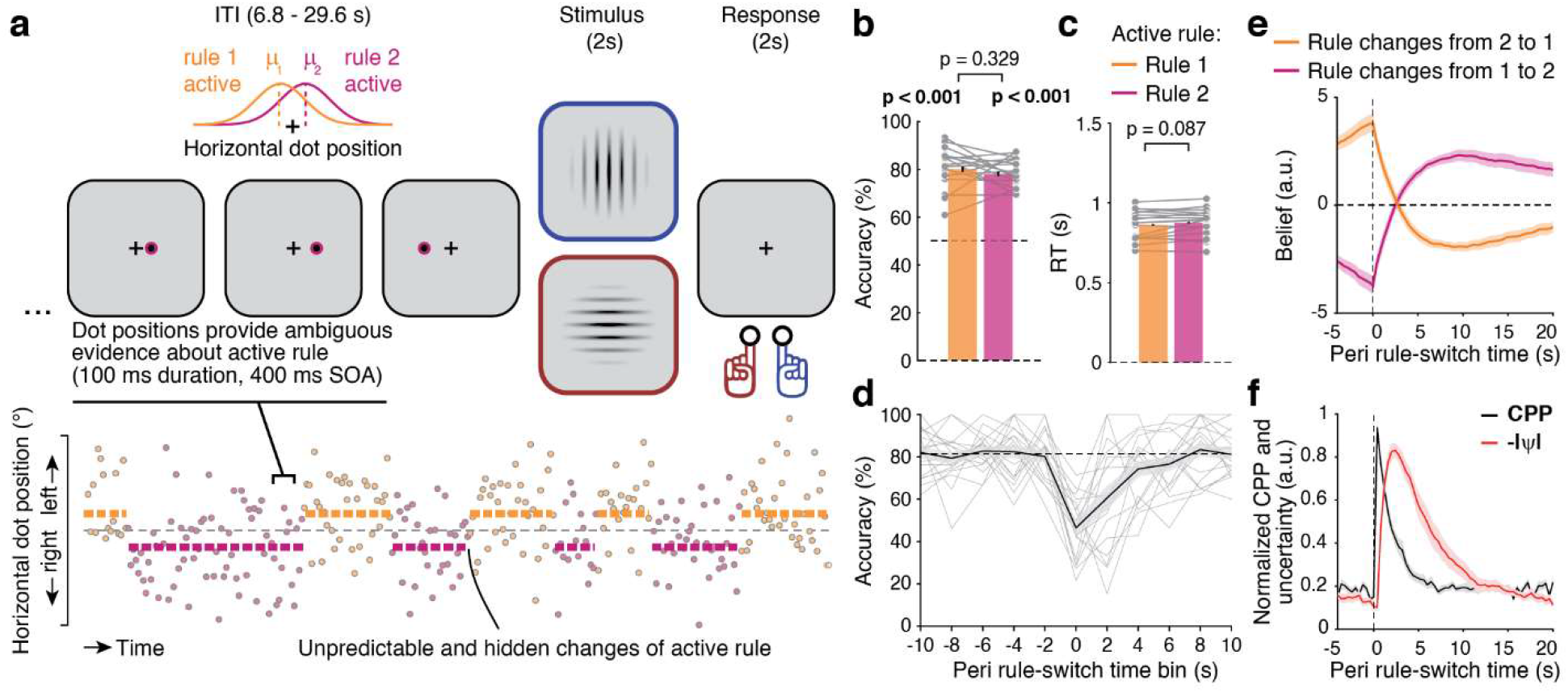
Task and behavior. **a**. Inferred rule task. Top: Example sequence of three evidence samples during the inter-trial-interval (ITI) preceding a task trial. Evidence samples were the horizontal positions of dots presented every 400 ms. Samples were drawn from one of two overlapping distributions, producing a noisy evidence stream. Bottom: example of noisy evidence stream. The generative distribution governed the active rule and could switch unpredictably between any two samples (probability: 1/70). **b**. Accuracy, and **c**. response time (RT) for each rule. The horizontal dotted line in b. shows chance level accuracy. Gray dots, individual participants. Bars, group average. Error bars, SEM. **d**. Accuracy for time bins centered on the onset of hidden rule changes. Accuracy drops substantially following a rule change and gradually recovers as participants accumulate evidence in favor of the rule change. Gray lines, individual participants. Black line, group average. Horizontal dotted line, average accuracy before the rue switch. Error bar, SEM. **e**. The belief-parameter from the model locked to rule switches. **f**. Model-derived behavioral variables locked to the onset of rule switches. Following a switch in rule, change point probability (CPP) and uncertainty (-|ψ|) increase. Error bar, SEM.

Participants were able to determine and apply the active rule well above chance level, and similarly across the two rules (Figure 2b,c; Table S1). Switches of the active rule resulted in a marked dip in performance, which then recovered over time as more sensory evidence in favor of the new rule was provided (Figure 2d).

The normative (Bayesian) strategy for solving this online inference task entails an adaptive, non-linear accumulation of evidence that balances build-up of stable belief states with sensitivity to the hidden (rule) switches (Glaze et al., 2015). We fit a model implementing this Bayesian inference strategy to participants’ choices (Methods). Adaptive sensitivity to rule switches was implemented in this model by a non-linear transformation of a prior before integrating it with newly arriving sensory evidence. The shape of this non-linear transformation depended on a subjective estimate of the probability of rule switches. The fitted model quantified the ‘belief’ (L) of the participant in favor of one rule over the other. This belief systematically changed sign after rule switches (Figure 2e).

Here, we focus on two other latent variables of the model that have been shown to modulate behavioral evidence accumulation profiles and pupil-linked arousal responses during active inference (Murphy et al., 2021): change point probability (CPP), and uncertainty (-|ψ|). CPP indicated the probability that a switch of rule has just occurred, given the participants’ subjective estimate of the probability of rule change, the new sensory evidence, and current belief. CPP was high in cases where a new sample of evidence was inconsistent with the participants’ current belief – in particular, when both the previous belief and new (contradictory) evidence were strong. Uncertainty about the environmental state (i.e., active SR rule) before encountering a new sample of sensory evidence was tracked by -|ψ|. Switches of active rule were followed by a peak in both quantities, with a rapid rise in CPP, and a subsequent and more protracted rise in uncertainty (Figure 2f).

In sum, hidden rule switches elicited a sequence of latent processes that included the detection of sensory evidence that violated previously held beliefs (CPP), a rise in uncertainty about the active rule (-|ψ|), and a flip in the sign of the belief state L. Ultimately, these internal events led to an adjustment of behavior to the new rule and, consequently, a recovery of task accuracy.

### Correlated variability of stimulus and action codes reflects active rule

Our prior work established that the variability of cortical population codes for stimulus and for action covaried, with a sign that depended on the model-derived belief of the participant (van den Brink et al., 2023). In the current study, we investigated the dependence of this correlated variability of cortical stimulus and action codes on brainstem arousal systems and the associated pupil dynamics. As a prerequisite for our analyses of the brainstem and pupil dynamics in the task (reported below), we first verified that the correlated variability also reflected the truly active rule, which did not always match participants’ internal belief about the active rule.

As in our original study (van den Brink et al., 2023), we quantified ongoing fluctuations of population codes as the graded output of decoders that captured stimulus orientation (horizontal vs. vertical) or chosen action (left hand vs. right hand response) discriminant patterns. These decoders were applied to the patterns of spontaneous fMRI signal fluctuations within a set of cortical and sub-cortical regions (Figure 3a; Table 1) from which stimulus and action-evoked activity had been removed. The resulting time series indicated if the activity pattern at any given moment tended toward vertical or horizontal stimulus orientation (for visual cortical regions), or towards left hand or right hand button press (for sub-cortical and cortical motor-related regions). We then correlated the stimulus and action decoder outputs separately for intervals corresponding to the two rules (Figure 3b,c).

**Figure 3.**
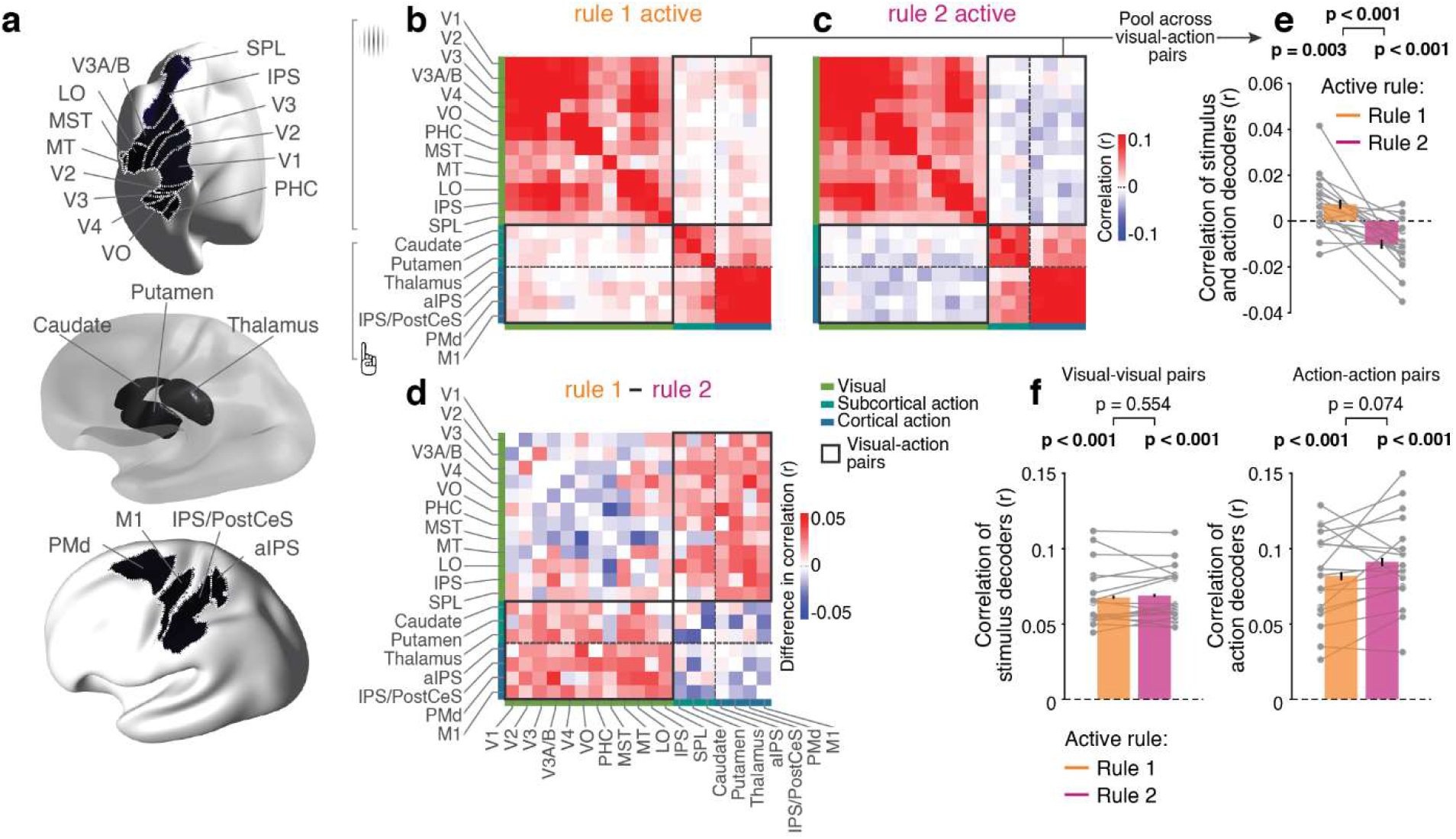
Correlated variability of population codes for stimulus and action. **a**. Regions of interest for the analysis of correlated variability. **b**., **c**., **d**., Correlation matrices for all stimulus and action decoders during rule 1 (b.), rule 2 (c.), and the difference between rules (d). Gray rectangles, stimulus-action pairs. **e**. correlations from (a.) and (b.), collapsed across all pairs in the gray rectangles. **f**. Correlations from (a.) and (b.), collapsed across all visual-visual pairs and action-action pairs (excluding values on diagonals). Gray lines, individual participants; bars, group average; error bars, SEM.

**Table 1.**
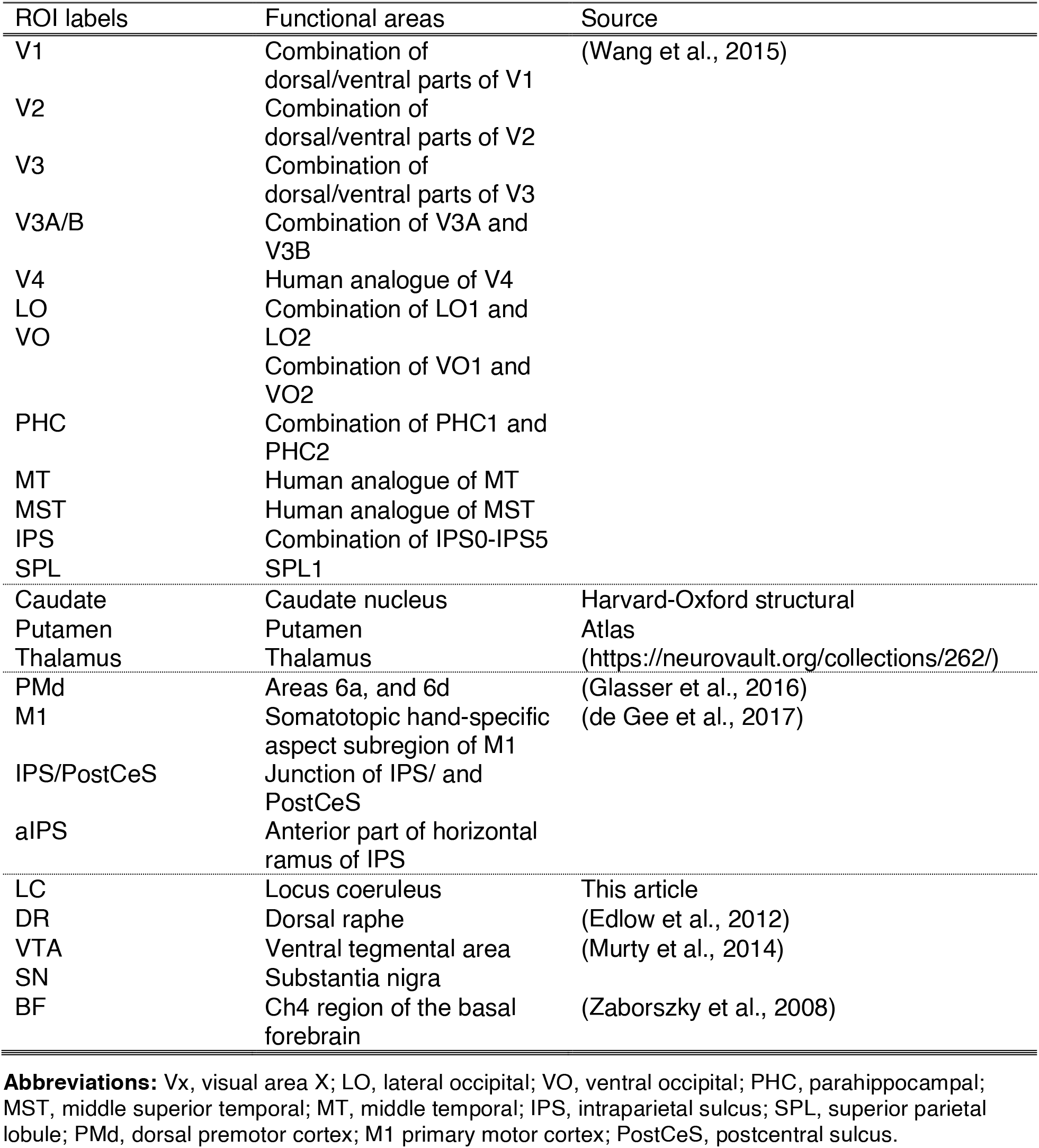
ROI definition.

The sign of correlation of the stimulus-action decoder pairs was a product of the design of the decoders: positive for coupling consistent with rule 1, and negative for rule 2 (cf. van den Brink et al., 2023). In line with the notion that the correlated variability depended on the active rule, the sign of correlations of stimulus-action decoder pairs differed between rules (Figure 3d,e; Table S1), while no such sign flip was observed for the correlations of stimulus-stimulus or action-action decoder pairs (Figure 3f; Table S1). We note that the active rule and model-inferred belief were strongly correlated, and we re-analyzed the data from our previous work. Hence, the results shown in Figure 3 were strongly expected based on the results from van den Brink et al (2022). They served merely as verification for our subsequent analyses of brainstem and pupil dynamics locked to the switches of the active rule.

### Brainstem neuromodulatory centers and pupil-linked arousal are recruited by rule switches

We next examined if rule switches also evoked activity in brainstem centers that release certain neuromodulators. We focused on five neuromodulatory centers (Figure 4a; Table 1): the cholinergic basal forebrain (BF), dopaminergic substantia nigra (SN) and ventral tegmental area (VTA), the serotonergic dorsal raphe (DR), and noradrenergic locus coeruleus (LC) (van den Brink et al., 2019). Imaging of the brainstem is difficult due to the size and location of the nuclei involved. We thus applied physiological noise correction and controlled for additional noise contained in ventricular signals (Brooks et al., 2013; de Gee et al., 2017). For optimal precision in localization, we delineated the smallest nucleus, the LC, within each individual, based on neuromelanin sensitive anatomical scans (Keren et al., 2009; de Gee et al., 2017) that highlighted the LC as hyperintense spots (Figure S1). The location of the resulting masks corresponded well to the known location of the LC in the posterior part of the pons, at the base of the fourth ventricle (Figure S2a). For the other (larger) regions, we relied on publicly available atlases (Table 1).

**Figure 4.**
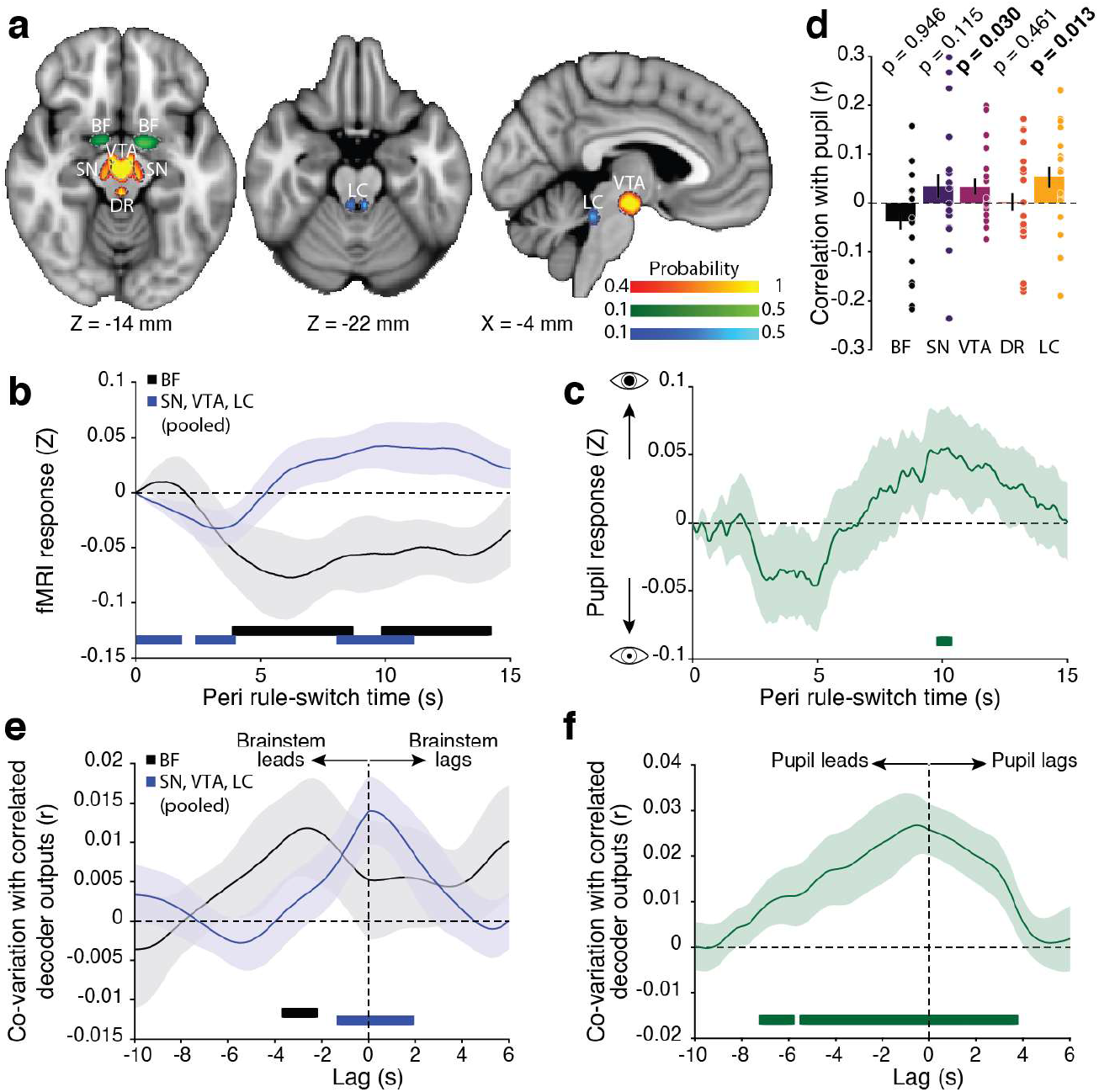
Relationship between brainstem neuromodulation, pupil dynamics, and correlated variability. **a**. Regions of interest within the brainstem, defined via neuromelanin sensitive scans of individual participants (LC), and anatomical atlases (other nuclei; Table 1). BF, basal forebrain. SN, substantia nigra. VTA, ventral tegmental area. DR, dorsal raphe. LC, locus coeruleus. **b**. Deconvolved response in the brainstem evoked by rule switches. **c**. Deconvolved pupil response evoked by rule switches. **d**. Covariation between the rule switch-evoked response magnitude between the brainstem and pupil. **e**. Cross-correlation between brainstem activity and stimulus-action decoder coupling within the cortex. Negative lags indicate activity in the brainstem preceding increases in decoder coupling. **f**. Cross-correlation between pupil diameter and stimulus-action decoder coupling within the cortex, corrected for hemodynamic delays. Negative lags indicate pupil dilation preceding increases in decoder coupling. In all panels, horizontal bars indicate *p* < 0.05, corrected for multiple comparisons with cluster-based permutation testing. Error bars, SEM.

Switches of the active rule caused prominent evoked activity in the catecholaminergic (i.e., dopaminergic and noradrenergic) nuclei (SN, VTA, and LC; Figure 4b; Table S1). These nuclei are strongly inter-connected (Sara, 2009) and have been implicated in orchestrating resets of cortical activity in response to contextual changes (Dayan & Yu, 2006; Fusi et al, 2007). We thus combined these three catecholaminergic brainstem nuclei in our analyses (Figure S2b shows individual nuclei). The DR did not show significant responses to rule switches (Figure S2b). Unexpectedly, the BF responded to rule switch events with a marked suppression of activity (Figure 4b; Table S1).

Non-luminance mediated fluctuations of pupil diameter are known to covary with activity in neuromodulatory centers of the brainstem, including the catecholaminergic nuclei (Murphy et al., 2014; McGinley et al., 2015a; Joshi et al., 2016; Reimer et al., 2016; de Gee et al., 2017; Larsen and Waters, 2018; Breton-Provencher and Sur, 2019). Indeed, we found pupil diameter to covary with the majority of nuclei as well (Figure S2c; Table S1). Critically, the pupil also dilated in response to the rule switches (Figure 4c; Table S1), and in a manner that was correlated with the rule-switch response of the VTA and LC (Figure 4d; Table S1).

In sum, hidden rule switches recruited catecholaminergic brainstem nuclei and pupil-linked arousal in a coordinated fashion. This is consistent with the idea that brainstem arousal systems may orchestrate the coupling of population codes for stimulus and action within the cortex.

### Brainstem and pupil fluctuations precede fluctuations of correlated variability

If brainstem arousal systems affect SR-coupling within the cortex, we expect to see increased activity in the brainstem prior to increases in cortical signatures of SR coupling strength. We thus cross-correlated the signals from all brainstem nuclei that responded to rule switches and covaried with the pupil (i.e. excluding the DR) and the pupil signal itself with the strength of stimulus-action decoder correlation. To this end, we evaluated the latter in a time-variant fashion. We used strength (i.e., absolute value) of the decoder correlations rather than the correlations per se, because we did not expect the brainstem signals to bias the decoder correlations in any particular direction (i.e., favor a specific SR rule), but rather to boost whichever coupling currently dominated (e.g., through plasticity mechanisms (Fusi et al., 2007)). We evaluated the correlations across a series of lags in order to chart temporal dependencies.

We found a pronounced peak at negative lags in the cross-correlation spectrum between BF and stimulus-action decoder coupling (Figure 4e; Table S1). This indicated that peaks of activity in the BF preceded peaks in stimulus-action decoder coupling. We also found significant correlations between activity of the pooled catecholaminergic nuclei and stimulus-action decoder coupling, including at negative lags, and peaking around zero lag (Figure 4e; Table S1). Finally, the pupil showed dilation before subsequent peaks in stimulus-action decoder coupling strength (Figure 4f; Table S1). These findings, specifically the lags of the correlations, could not be explained by hemodynamic or pupillary delays because all delays were accounted for in the analysis. Moreover, because the pupil signal was sampled at a higher rate than the brainstem signal, it may be more sensitive to short (∼1s) lags between brainstem responses and changes in SR-coupling within the cortex.

Together, our results are consistent with the idea that activity in neuromodulatory nuclei (within the BF, and indexed by pupil diameter) tracked hidden rule switches, and modulated rule-specific coupling of stimulus and action codes amongst the network of brain regions that implemented the primary decision process. The brainstem receives input from brain areas that are involved in active inference, such as anterior cingulate (Aston-Jones and Cohen, 2005). We thus next asked if the arousal system encoded latent variables of behavior that are informative of rule switches.

### Pupil dynamics encode computational variables involved in rule inference

Sensitivity of the arousal system to rule switches may be conferred by latent variables that track changes in environmental state: CPP and uncertainty (-|ψ|) (Nassar et al., 2012; Murphy et al., 2021). In our task, samples of evidence were presented without temporal jitter and at a rapid pace (Figure 2a), likely too quickly to relate brainstem fMRI responses to sample-wise estimates of CPP and -|ψ|. Nevertheless, the pupil was sampled at a faster rate than the fMRI signal.

Given its close correspondence to brainstem activity, both expected from prior work (Murphy et al., 2014; McGinley et al., 2015a; Joshi et al., 2016; Reimer et al., 2016; de Gee et al., 2017; Larsen and Waters, 2018; Breton-Provencher and Sur, 2019), and observed here (Figure 4, Figure S2c), we used the pupil as a summary signal for probing relationships with computational variables of behavior.

CPP, a latent variable in a model fit to participants’ choices, increased rapidly following rule switches (Figure 2f). The temporal derivative of pupil diameter is maximally sensitive to such rapid changes (Murphy et al., 2021), and closely tracks activity in the noradrenergic LC (McGinley et al., 2015a; Reimer et al., 2016). We thus regressed CPP onto the derivative of diameter, while controlling for a range of other variables. Shortly after evidence sample onset, the derivative of diameter indeed encoded CPP (Figure 5a; Table S1). Uncertainty evolved more slowly than CPP (Figure 2f). We therefore expected its signature to become visible in diameter (rather than its derivative). This is indeed what we found (Figure 5b; Table S1). In conclusion, the pupil, a signal that closely matched the dynamics of brainstem neuromodulatory nuclei, tracked the two latent computational variables of the inference process that were sensitive to changes in environmental state (i.e., SR rule).

**Figure 5.**
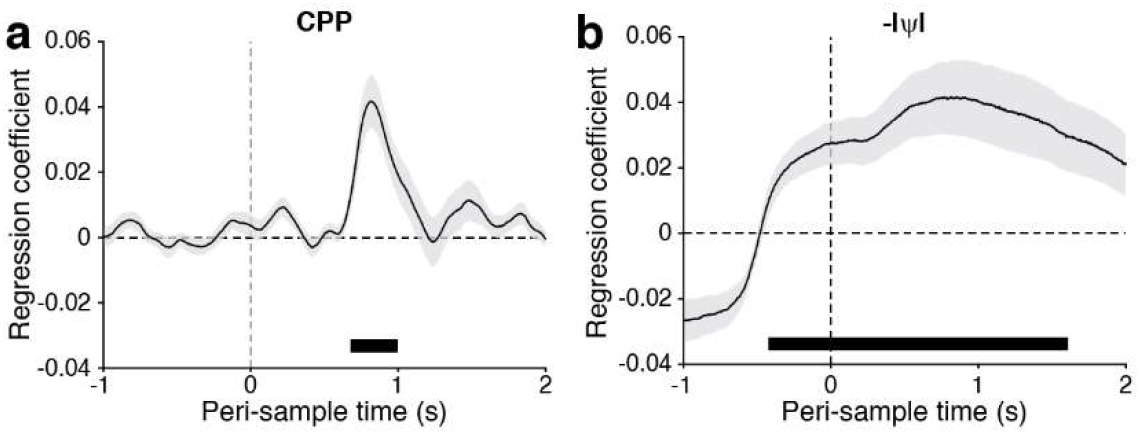
Relationship between pupil dynamics and behavior. **a**. Relationship between the derivative of pupil diameter and change point probability (CPP). **b**. Relationship between pupil diameter and uncertainty. In all panels, horizontal bars indicate *p* < 0.05, corrected for multiple comparisons with FDR correction. Error bars, SEM.

## Discussion

It has long been established that neuromodulatory systems shape the global state of, and plasticity within, cortical networks. Much less is known about whether and how the interplay between the brainstem and cortex shapes the neural bases of specific cognitive behaviors – such as context-dependent sensory-motor decisions that we studied here. We found that unpredictable switches in the state of the environment (i.e., SR rule) triggered a boost of arousal: a response within the centers that release neuromodulators, which coincided with a dilation of the pupil. Pupil-linked arousal also encoded rapid fluctuations of key computational variables for detecting hidden rule switches. Finally, boosts in the activity of brainstem arousal systems were followed by stronger coupling of stimulus and action selective patterns within the cortical regions that implement decisions. Taken together, our results are consistent with the idea that brainstem arousal plays an important role in shaping large-scale cortical pathways for flexible cognitive behavior in uncertain environments.

A key aspect of adaptive behavior is that SR-associations must be able to be formed in an arbitrary manner. A number of previous studies on flexible decision-making have been done with non-arbitrary SR-associations (Heinzle et al., 2012; Sarafyazd and Jazayeri, 2019; Duan et al., 2021), where a stimulus is associated with a prepotent response that must be conditionally acted upon or suppressed, such as pro- and anti-saccade tasks (Munoz and Everling, 2004). By contrast, SR-associations in our task were inherently symmetrical, where neither stimulus had a prepotent response. Therefore, default anatomical pathways that establish SR-mapping within the cortex are unlikely be in place before SR-associations are learned. These arbitrary SR-associations could come about through plasticity mechanisms that reshape existing SR-pathways or form new ones when no default pathways exist. This notion finds support in work in non-human primates and computational modeling, which has suggested that flexible SR-associations are the result of plasticity mechanisms operating on multiple time scales (Fusi et al., 2007).

Our current findings align well with this mechanistic interpretation. In this scenario, synaptic plasticity mechanisms reshape the pathways along which sensory information is directed to motor cortex (Fusi et al., 2007), prompted by neuromodulator release in the cortex in response to sensory evidence in favor of a new active rule. Neuromodulator release elicits plasticity of cortical (Bear and Singer, 1986; Rasmusson, 2000; Berridge and Waterhouse, 2003; Huang et al., 2004; Vetencourt et al., 2008; Marzo et al., 2009; Nadim and Bucher, 2014) and striatal connections involved in learned associations (Reynolds et al., 2001; Reynolds and Wickens, 2002). Moreover, neuromodulator release occurs in response to behaviorally relevant stimulus features (Berridge and Waterhouse, 2003; Aston-Jones and Cohen, 2005; Bouret and Sara, 2005; Sarter et al., 2009). Neuromodulators are thus well suited to restructure SR-association pathways when dictated by the state of the environment. Our findings that arousal responses followed in reaction to rule switches, but preceded the cortical signatures of remapped SR-associations, are consistent with this scenario. Our findings also align with theoretical proposals that link phasic arousal signals to cortical reorganization in response to changes of environmental state (Bouret and Sara, 2005; Dayan and Yu, 2006)

One prominent line of work focuses on the role of frontal and parietal associative regions in flexible SR mapping. Such regions have been found to encode rule information (Miller and Cohen, 2001; Woolgar et al., 2016; van den Brink et al., 2023), and they have been proposed to flexibly route information from sensory to motor cortices (Miller and Cohen, 2001). These regions may do so by transiently activating in response to the conjunction of stimulus and rule information and acting as switches between the sensory and motor cortices (Cocuzza et al., 2020; Kikumoto and Mayr, 2020; Ito et al., 2022). Other findings indicate that prefrontal cortex encodes key computational variables for performance monitoring and tracking environmental state (Behrens et al., 2007; O’Reilly et al., 2012; McGuire et al., 2014). Frontal and parietal regions are known to innervate the brainstem arousal system (Schwarz and Luo, 2015; Schwarz et al., 2015; Breton-Provencher and Sur, 2019). Our present findings thus raise the possibility that prefrontal association cortex may control flexible SR information flow indirectly, through conveying computational variables such as change-point probability and uncertainty to the brainstem, which in turn sculpt the neural SR pathways accordingly.

The negative response to rule switches that we observed in the BF (Figure 4b) was unexpected, but interesting in light of recent findings. Positive fMRI transients across wide areas of the cortex, and especially in sensory cortices, have been shown to co-occur with drops of activity in the BF region (Liu et al., 2018). In addition, in non-human primates inactivation of the BF results in suppression of correlated fMRI signal fluctuations that are topographically aligned with its afferents (Turchi et al., 2018). Notably, although this region is the main source of cortical acetylcholine (Mesulam and van Hoesen, 1976; Mesulam et al., 1983; Mesulam and Changiz, 1988), it is diverse and also sends prominent long-range GABAergic projections to the cortex (Lin et al., 2015). Thus, it is possible that the here observed suppression of BF activity following rule switches represented a disinhibitory signal (Letzkus et al., 2015) that allowed correlated fluctuations in the cortex to emerge. This possibility is particularly appealing considering the proposed role of disinhibitory signals in flexible information routing within the cortex (Wang and Yang, 2018). Nevertheless, our findings also indicated that increases in BF activity preceded increases in SR-coupling (Figure 4e). It is thus also possible that a suppression of BF activity elicited by rule switches triggered a suppressive effect on SR-coupling within the cortex, and a new instantiation of SR-coupling was brought about by the subsequent activity in the catecholaminergic nuclei. Causal manipulations of specific neuron types within the BF, and concurrent electrophysiological recordings within the cortex, would be well suited to arbitrate between these alternatives.

Due to the nature of the fMRI signal, we cannot definitively distinguish between neuromodulatory signals and those signals originating from other sources. In addition, imaging of nuclei within the brainstem in general is difficult due to their size and proximity to noise sources (Brooks et al., 2013). We have used several methodological approaches to mitigate these issues: by optimizing slice alignment with respect to the brainstem and correcting for physiological noise. We also created masks for the LC on an individual participant basis for optimal localization (Eckert et al., 2010). Finally, we confirmed the expected covariation between signals extracted from individual nuclei and pupillary indices of arousal, for both ongoing signal fluctuations and the more specific rule-evoked responses (Figure S2c, Figure 4d). Together, these choices in experiment design, analysis, and findings, increased the likelihood that the signals we measured are truly related to neuromodulatory nuclei.

Pupil diameter covaried with uncertainty, which was significant before the onset of a new sample of evidence (Figure 5b). Although part of this pre-cue onset covariation can be accounted for by the fact that the prior is the result of sensory evidence that precedes the current sample, we cannot fully exclude that part of this effect is driven by correlation in the values of uncertainty at adjacent cues. In future studies this can be ruled out by using designs with larger lags and jitter between consecutive cues so that protracted pupil responses to individual cues are more clearly separable. Nonetheless, the key finding that uncertainty covaries with pupil diameter holds regardless of the precise timing with respect to the evidence samples.

In conclusion, during flexible decision-making unpredictable switches in environmental state engage the arousal system. The arousal response in turn is followed by coordinated shifts of coupling between stimulus and action specific activity within the cortical regions that implemented the decision process. Thus, brainstem arousal may orchestrate the association of stimuli with their appropriate actions.

## Acknowledgements

This work was funded by the Deutsche Forschungsgemeinschaft (DFG, German Research Foundation) – DO 1240/3-1 (THD), DO 1240/4-1 (THD), and SFB 936 - 178316478 - A6 (CB), A7 (THD) & Z3 (THD).

## Author contributions

**RLvdB:** Conceptualization, Methodology, Software, Formal analysis, Resources, Data Curation, Writing – Original Draft, Writing – Review & Editing, Visualization. **KH:** Methodology, Software, Formal analysis, Data Curation, Writing – Review & Editing. **NW:** Conceptualization, Methodology, Software, Formal analysis, Investigation, Data Curation, Writing – Review & Editing, Supervision. **PRM:** Methodology, Software, Formal analysis, Writing – Review & Editing. **JF:** Methodology, Writing – Review & Editing. **JC-T:** Methodology, Software, Formal analysis, Writing – Review & Editing. **CB:** Conceptualization, Writing – Review & Editing. **THD:** Conceptualization, Methodology, Resources, Writing – Review & Editing, Visualization, Supervision, Project administration, Funding acquisition.

## Materials and Methods

The current study involves the reanalysis of previously published data (although all presented findings are novel). For a full description of all task parameters and preprocessing, please see van den Brink et al. (2023).

### Participants

A total of 22 healthy individuals with normal or corrected vision (median age 27, range 21 – 44, 8 male) took part in our experiment. All participants gave written informed consent and the study was approved by the ethics committee of the Hamburg Medical Association. Four participants were excluded from the current study: one for failure to complete all three sessions, another three for technical reasons (failure to record physiological signals or response data). The final N was thus 18.

### Behavioral task

Participants performed two different versions of a flexible decision-making task, of which only one (the ‘inferred rule’ task) is analyzed in the current study. This task involved a basic visual orientation discrimination judgment (lower-order decision), combined with the selection of a volatile sensory-motor (SR) mapping rule (higher-order decision). The SR-mapping rule (Figure 1a) determined the correct action for a given orientation judgment (Figure 2a).

The active rule had to be continuously inferred from a sequence of noisy sensory evidence samples, which were small dots that appeared in rapid succession on the horizontal meridian (Figure 2a). The dots were drawn from one of two generative Gaussian distributions with equal SD and means that were equidistant from the central fixation point but on opposite sides. The generative distribution at any moment determined the active rule. Critically, the active rule could change unpredictably from one sample to the next, with a low probability (hazard rate) of 0.0143. Thus, in order to determine the active rule at a given moment, participants needed to continuously integrate the noisy rule evidence over time.

The appearance of the stimulus for the lower-order decision prompted participants to report their orientation judgement. ITIs for the lower-order decision were long and variable (uniform: 6.8 – 29.6 s). The accuracy of the action depended on both the selection of the correct rule, and on the correct orientation judgment.

### MRI data collection

MRI scans were conducted using a Siemens PrismaFit 3T MRI scanner with a 64-channel head-neck coil. We collected 6 runs of the inference task, split over two sessions (T2*-weighted EPI data; Flip angle: 70°; TR: 1.9 s; TE: 28 ms; FOV: 224 × 224 mm^2^, 62 slices (no gap) of 2.0 mm isotropic voxels; 328 volumes), and simultaneously recorded cardiac pulsation and breathing using a pulse oximeter and pneumatic belt. Slices were oriented perpendicular to the rostral-caudal axis of the brainstem as to maximize SNR in this region.

On both sessions, we collected B0 field homogeneity scans (Phase difference and magnitude image: flip angle: 40°; TR: 0.678 s; TE: 5.42/7.88 ms). At the end of the first MRI session, we collected a high resolution T1 neuromelanin sensitive scan, for the purpose of locating individual participants’ LC (T1-TSE; Flip angle: 120°; TR: 675 ms; TE: 12 ms; FOV: 175 × 224 mm^2^, 14 slices (2.0mm, no gap); 0.70x0.70 mm^2^ interpolated to 0.35x0.35 mm^2^). The slices were oriented perpendicular to the rostral-caudal axis of the brainstem in order to align with the longitudinal extent of the LC. At the end of the second MRI session, we collected a whole-brain T1 anatomical scan (MPRAGE; Flip angle: 9°; TR: 2.3 s; TE: 2.98 ms; TI 1.1 s; FOV: 192 × 256 mm^2^, 240 slices; 1.0 × 1.0 × 1.0 mm).

### Pupil recording and preprocessing

Pupil diameter and gaze position were recorded at 1000 Hz with an MRI-compatible EyeLink 1000 eye tracker and calibrated with a 9-point fixation routine. Blinks and missing data segments were linearly interpolated in the eye position data. Other artifacts were identified by the derivative of the pupil diameter exceeding a threshold of 25 pixels and interpolated across. This process was repeated iteratively to ensure all artifacts were identified and removed, in accordance with prior work (van den Brink et al., 2016). All sections containing artifacts were then similarly interpolated across in the gaze x and y position data.

### Behavioral modeling

#### Normative model

The full details of the normative model and fit are provided in van den Brink et al. (2023). In brief, the rule selection (the higher-order decision) process was cast as a dynamic belief updating process (Glaze et al., 2015). In this model, a belief *L* for one possible state (rule 1) versus the alternative possible state (rule 2) was computed for each sample of evidence. This belief then underwent a non-linear transformation, dependent on a subjective hazard rate *Ĥ*, in order to become a prior (*ψ*) on the presentation of the next sample, thus rendering the model adaptive to volatility of the rule.

#### Model-derived uncertainty and change-point probability

We used the model fits to produce two key computational parameters associated with each evidence sample: Uncertainty, and change-point probability (CPP). Uncertainty associated with the selected rule was computed as the negative of the magnitude of the prior ψ. CPP is a computational quantity that both the normative belief updating process and human participants are sensitive to in volatile decision-making contexts such as ours (Murphy et al., 2021). CPP was the posterior probability of a change having occurred, given *Ĥ*, the previous posterior and the new evidence sample. CPP was computed as follows:

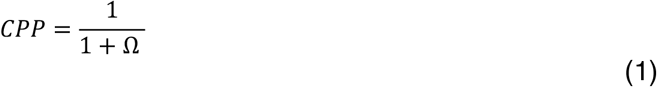

where,

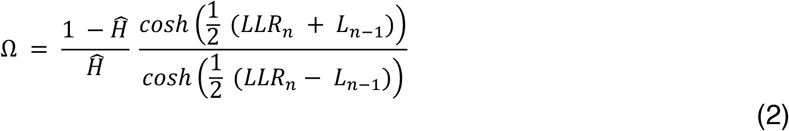

and where *n* was the current sample, LLR was the sensory evidence in favor of one rule over the other in the form of a log likelihood ratio. Eq. 1 and 2 represent an algebraic rearrangement of the equation for CPP derived by Murphy et al. (2021).

### MRI data analysis

#### Pre-processing of fMRI data

The fMRI data were preprocessed identically to van den Brink et al. (2023). In brief, preprocessing included: motion correction, skull stripping, B0 unwarping, high-pass filtering at 100s, prewhitening, physiological noise correction with retrospective image correction (RETROICOR) (Glover et al., 2000), slice-time correction, and non-linear registration to MNI space.

No spatial smoothing was applied to the fMRI data for two reasons: i) To preserve spatial patterns that were the basis of multi-voxel pattern analysis (see below), and ii) to prevent signal mixing between the brainstem and closely adjacent 4^th^ ventricle (Brooks et al., 2013).

#### Delineation of ROIs

Regions of interest (ROIs) were defined using a variety of sources (Table 1), including publicly available atlases as well as individually created masks based on neuromelanin sensitive scans. The set of ROIs that we used for all SR coupling-related analyses (see below) presented in this article were selected to span the sensory-motor pathway underlying the lower-order decision (Table 1; Figure 3a). Because the probabilistic masks of neuromodulatory brainstem centers (Zaborszky et al., 2008; Edlow et al., 2012; Murty et al., 2014) varied substantially in their spatial extent, we confined each of these masks to 18 peak probability voxels. Masks of comparable size for the LC were defined for individual participants based on neuromelanin sensitive TSE scans (see below).

#### Individual delineation of LC ROIs

Masks of the LC for individual participants were created using a semi-automated procedure. First, a mask was manually drawn on the approximate locations of bilateral LC in the T1-TSE scans (Sasaki et al., 2006; Keren et al., 2009), which coincided with the known anatomical location of the LC within the pons, along the floor of the fourth ventricle. An automated algorithm then identified the peak intensity voxel within the manually drawn mask, and an additional 14 contiguous voxels with the highest intensity in each hemisphere. Individual masks (Figure S1) were co-registered to 2 mm isotropic MNI space together with the high-resolution whole-brain T1 scans, max-unit normalized, and thresholded at values of 0.1 to reduce partial voluming effects.

A probabilistic map for the group was defined as the proportion of participants with non-zero values for each voxel in MNI space. This group mask was compared to a previously published mask of the LC (Keren et al., 2009) in order to verify the average location (Figure S2a).

#### Brainstem time-series extraction and post-processing

Imaging of the brainstem is difficult due to the size of the nuclei involved and their proximity to noise sources such as the 4^th^ ventricle (Brooks et al., 2013; Turker et al., 2021). Physiological noise in the functional data was mitigated using slice-specific RETROICOR (see: van den Brink et al. (2023)). Nevertheless, in order to maximize the anatomical specificity of the extracted signals, we weighted the time-series of each brainstem nucleus by the probability of voxels within the respective masks. Furthermore, we extracted a probability-weighted time series of voxels within a mask of the 4^th^ ventricle. Variance associated with the ventricular signal was then removed from the signals of the individual brainstem neuromodulatory nuclei using linear regression.

#### Time-resolved correlation of stimulus and action population codes

We quantified the coupling between population codes for stimulus and action by means of time-resolved decoder output correlation (Anzellotti and Coutanche, 2018), following the methods described in van den Brink et al. (2023). Specifically, we trained sets of support vector machine (SVM) classifiers to decode stimuli (horizontal versus vertical orientation) and action (left versus right hand button press) based on evoked response patterns within cortical and subcortical regions (Figure 3a) that spanned the sensorimotor network.

We then removed feature-specific evoked responses that the decoders were trained on from the data using deconvolution, thus isolating ongoing fluctuations (i.e. activity pattern fluctuations that were not driven by choice grating onset or button presses). We next projected the decision functions (weight vectors) from the stimulus and action decoders onto these estimates of ongoing multi-voxel activity patterns. This produced vectors that compactly summarized ongoing pattern fluctuations for stimulus (*S*; for visual cortical regions), or action (*A*; for motor cortical and subcortical regions). For example, in V1, the time course *S* reflected the extent to which patterns of ongoing activity spontaneously tended towards vertical or horizontal orientations (Kenet et al., 2003).

The sign of *S* and *A* resulted from the specific decoder design: positive values indicated an ongoing activity pattern that resembled responses to vertical gratings (or left hand button presses) and negative values indicated an ongoing activity pattern that resembled responses to horizontal gratings (or right hand button presses). The schematic in Figure 1b in combination with this decoder design non-trivially predicted a positive correlation of decoder outputs for rule 1 and a negative correlation for rule 2, for the stimulus-action pairs specifically.

We Z-scored *S* and *A* across time and computed, for each time point *t* in the fMRI time series, the degree of covariation between the stimulus and action patterns, via:

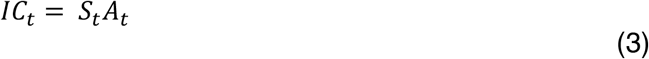

Note that because *S* and *A* were Z-scored, *IC*, if averaged along the time dimension, was identical to the Pearson correlation coefficient of *S* and *A*. Therefore, *IC* represented the correlation of stimulus and action population codes unwrapped along the time dimension.

We tested if *IC*, averaged across all visual-motor ROI pair combinations, differed between sections of data that were split according to the active rule, using non-parametric permutation testing (10,000 iterations).

#### Rule switch-evoked brainstem and pupil responses

We used deconvolution to test for the sensitivity of the brainstem and pupil to switch events in the active rule, without making any assumptions about the shape of the hemodynamic or pupillary response (Dale, 1999). Specifically, the estimated response *R* (of shape P x 1) was obtained via:

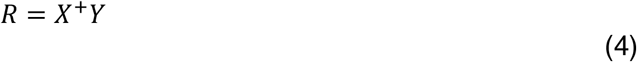

where *Y* was an *N* x 1 linearly detrended and z-scored time series (up-sampled fMRI data, or pupil diameter down-sampled to 50 Hz). *X* was an *N* x *P* design matrix, with value 1 in every element along those diagonals corresponding to the occurrence of switches in active rule, ^+^ denoted the pseudoinverse, and *P* was peri-stimulus time.

The deconvolved responses were compared to chance (zero) and corrected for multiple comparisons across time points using non-parametric cluster-based permutation testing (10,000 iterations, two-tailed) (Maris and Oostenveld, 2007).

#### Correlation between brainstem activity and pupil diameter

We assessed the correlation between brainstem and pupil activity in two ways: i) by correlating estimates of single-switch responses in the brainstem and in the pupil (Figure 4c), and ii) by correlation of the entire brainstem activity and pupil time series (Figure S2c).

Single-switch response amplitude scalars for individual brainstem nuclei were computed by regression of the individually estimated HRF (see section: *Rule switch-evoked brainstem and pupil responses*) onto the single rule switch locked data. Similarly, we computed single-switch responses in the pupil via regression, using the standard pupillary response function described by Hoeks and Levelt (1993), with a peak latency set for individual participants according to the deconvolved pupil response. The resulting vectors of response amplitude scalars were then correlated to estimate covariation in evoked responses in the brainstem and pupil.

Whole time-series correlation (Figure S2c) between the brainstem signals and pupil signal was done including a correction for HRF and pupillary response delays. That is, we estimated a peak HRF latency for individual participants from the stimulus-evoked response in V1 (van den Brink et al., 2023), and shifted the pupil signal forward by this latency. Because the pupil signal itself also lags neural activity, we subsequently shifted the pupil signal back by 1 s, following prior work (Yellin et al., 2015).

The significance of both single-switch response covariation as well as covariation of the full time series was assessed with non-parametric permutation testing (10,000 iterations). We tested one-tailed because we expected positive covariation of activity in the pupil and brainstem.

#### Cross-correlation between stimulus-action code covariation, brainstem activity, and pupil

We used cross-correlation to examine temporal relationships between stimulus-action code covariation (computed via Eq. 3) and two other signals: activity in the brainstem, and pupil diameter. We used cross-correlation because the temporal sequence of activity fluctuations may be indicative of directional relationships between these signals. That is, if the activity in the brainstem modulates coupling at the level of the cortex, then we expect the brainstem to become active prior to an increase in cortical coupling.

Stimulus-action code covariation was a signed signal, where the sign depended on the active task rule. We expected the brainstem to modulate the magnitude of this signal, regardless of sign, so in all related analyses we used the absolute of stimulus-action code covariation.

Because the pupil signal lags putative neural activity that underlies it, we corrected for this delay by shifting the pupil signal backward by the expected latency of 1s (Yellin et al., 2015), and aligned it with the hemodynamic signal by shifting forward by the observed hemodynamic delay (identically to the procedure described in the section: *Correlation between brainstem activity and pupil diameter*). Stimulus-action code covariation and brainstem activity were both hemodynamic signals in origin, so no lag correction was required before cross-correlation.

Correlation coefficients of the cross-correlations were compared to chance (zero) and corrected for multiple comparisons across time points using non-parametric cluster-based permutation testing (10,000 iterations, two-tailed) (Maris and Oostenveld, 2007).

#### Relationship between pupil dynamics and model computational variables

We examined the relationship between pupil and behavior with linear regression of the variables resulting from the computational model fit to individual participants’ behavior onto (the derivative of) pupil diameter. The regression had the following form:

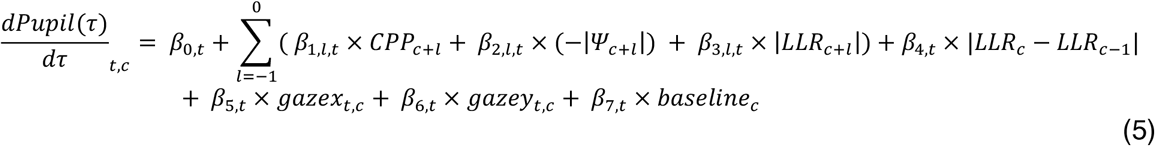

where *t* was time relative to cue (evidence sample) onset, *c* indexed cues, 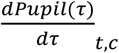 gave the value of the pupil derivative time series at time *t* relative to the onset of cue *c*. CPP was change point probability, -|Ψ| was uncertainty, LLR was the sensory evidence (i.e., the log-likelihood ratio), *gazex* and *gazey* were gaze positions on screen, and baseline was average pupil diameter in a 1 s window preceding cue onset. The terms relating to prior cues were included to account for auto-correlation in the pupil response and isolate responses to the current cue. The magnitude of the difference between the current and previous cue LLR was included to exclude that any relationship between pupil and CPP was driven by low-level visual differences between consecutive cues.

We also fit a variant of this regression model to baseline corrected pupil diameter (rather than the derivative). This regression model did not include baseline diameter, or terms for preceding cues:

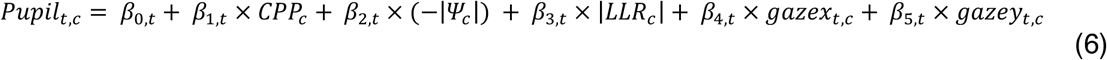

Based on prior work (Murphy et al., 2021), we expected CPP following cue onset to covary positively with the derivative of pupil diameter. We also expected uncertainty to covary positively with pupil diameter.

Regression coefficients were compared to chance (zero) using non-parametric permutation testing (10,000 iterations, one-tailed) and corrected for multiple comparisons using the false discovery rate (FDR).

## Supplementary Figures and Tables

**Figure S1.**
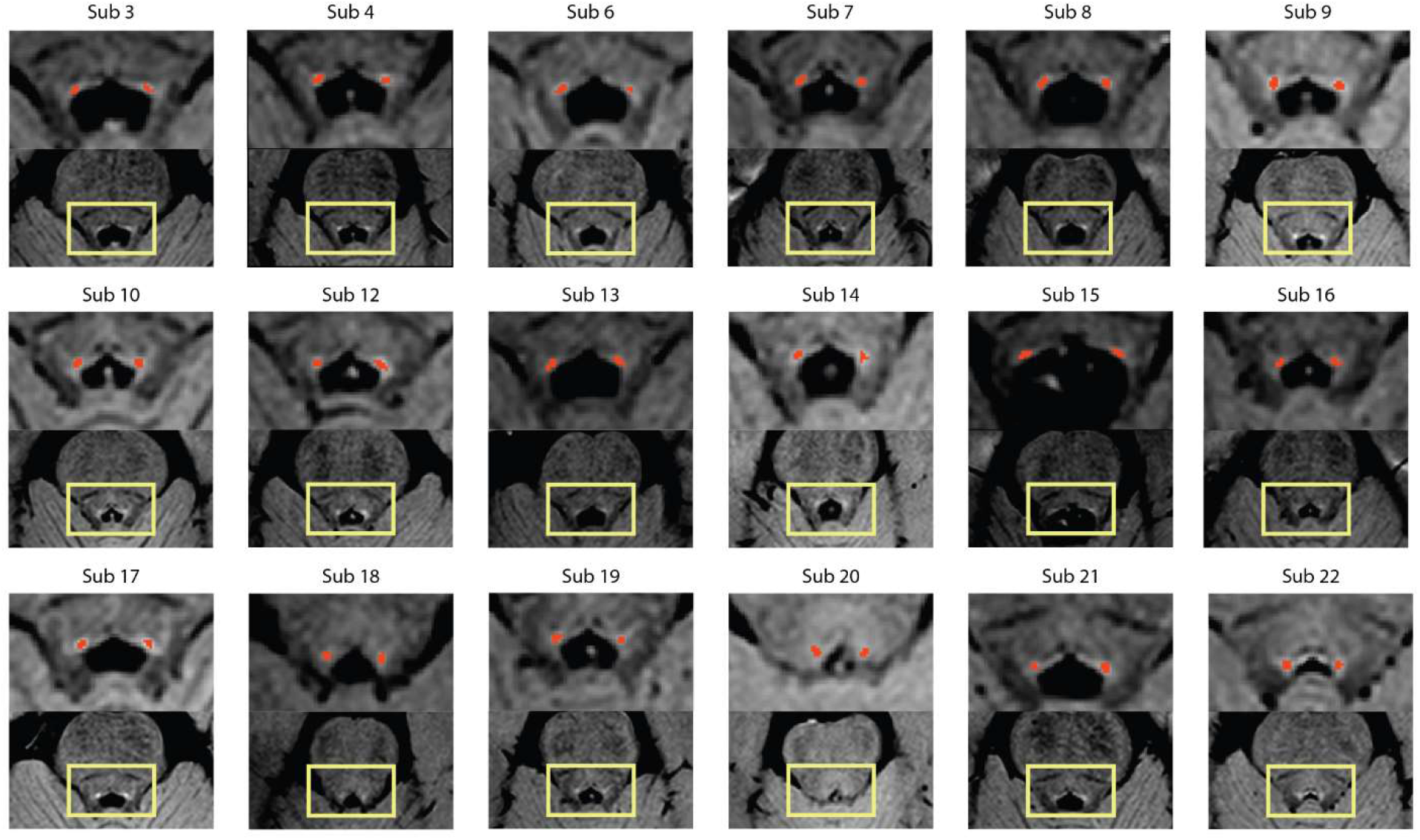
Neuromelanin sensitive TSE scans for all participants. The mask of the locus coeruleus (LC) is overlayed in red. The masks were created by manually selecting the maximal voxel within the LC region on each side, and subsequently automatically selecting 15 additional contiguous voxels with the highest intensity.

**Figure S2.**
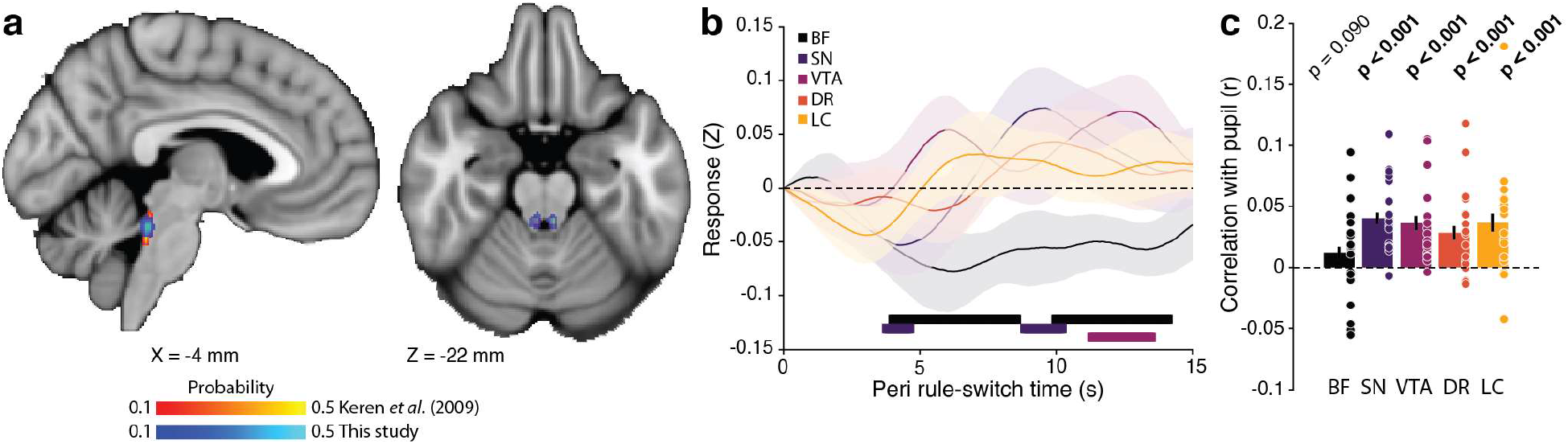
**a**. Overlap of LC mask of the current study with that of Keren *et al*., (2009). **b**. Deconvolved response in the brainstem evoked by rule switches, shown for all individual nuclei. **c**. Correlation between brainstem signals and pupil diameter, across time. Covariation between the rule switch-evoked response magnitude between the brainstem and pupil. Dots, individual participants. Bars, group average. Error bars, SEM. Horizontal bars indicate *p* < 0.05, corrected for multiple comparisons with cluster-based permutation testing.

**Table S1.**
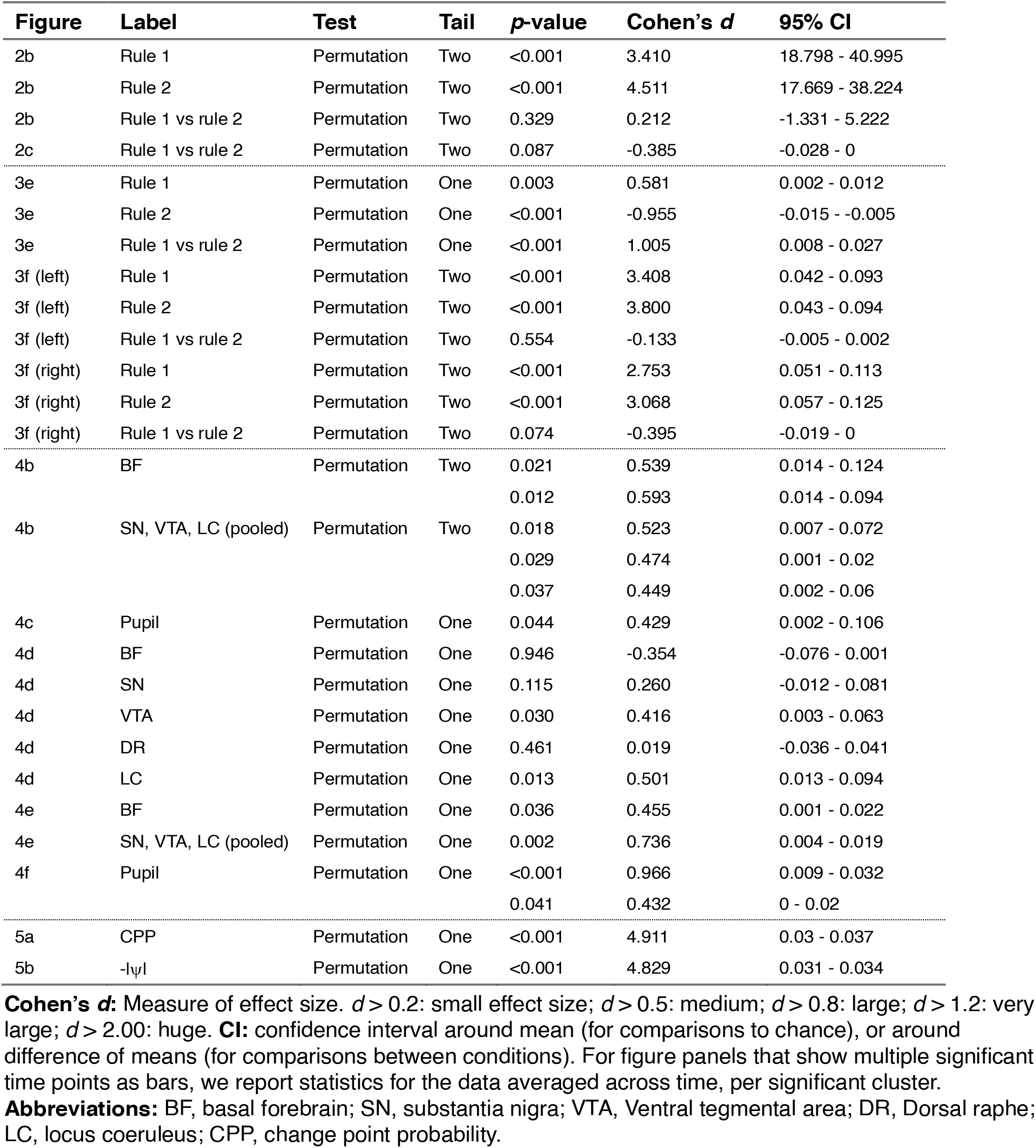
Statistics summary.

## References

Anzellotti S, Coutanche MN (2018) Beyond Functional Connectivity: Investigating Networks of Multivariate Representations. Trends in cognitive sciences 22:258–269.

Aston-Jones G, Cohen JD (2005) An integrative theory of locus coeruleus-norepinephrine function: Adaptive gain and optimal performance. Annu Rev Neurosci 28:403–450.

Bear M, Singer W (1986) Modulation of visual cortical plasticity by acetylcholine and noradrenaline. Nature 320:172–176.

Behrens TE, Woolrich MW, Walton ME, Rushworth MF (2007) Learning the value of information in an uncertain world. Nature neuroscience 10:1214–1221.

Berridge CW, Waterhouse BD (2003) The locus coeruleus–noradrenergic system: modulation of behavioral state and state-dependent cognitive processes. Brain Research Reviews 42:33–84.

Bogacz R, Brown E, Moehlis J, Holmes P, Cohen JD (2006) The physics of optimal decision making: a formal analysis of models of performance in two-alternative forced-choice tasks. Psychol Rev 113:700–765.

Bouret S, Sara SJ (2005) Network reset: a simplified overarching theory of locus coeruleus noradrenaline function. Trends Neurosci 28:574–582.

Breton-Provencher V, Sur M (2019) Active control of arousal by a locus coeruleus GABAergic circuit. Nature neuroscience 22:218–228.

Brooks JC, Faull OK, Pattinson KT, Jenkinson M (2013) Physiological noise in brainstem FMRI. Frontiers in human neuroscience 7:623.

Cocuzza CV, Ito T, Schultz D, Bassett DS, Cole MW (2020) Flexible Coordinator and Switcher Hubs for Adaptive Task Control. The Journal of neuroscience 40:6949–6968.

Dale AM (1999) Optimal experimental design for fMRI. Human Brain Mapping 8:109–114.

Dayan P, Yu AJ (2006) Phasic norepinephrine: a neural interrupt signal for unexpected events. Network: Comp in Neur Sys 17:335–350.

de Gee JW, Colizoli O, Kloosterman NA, Knapen T, Nieuwenhuis S, Donner T (2017) Dynamic modulation of decision biases by brainstem arousal systems. eLife 6:e23232.

Donner TH, Siegel M, Fries P, Engel AK (2009) Buildup of choice-predictive activity in human motor cortex during perceptual decision making. Current biology : CB 19:1581–1585.

Duan CA, Pagan M, Piet AT, Kopec CD, Akrami A, Riordan AJ, Erlich JC, Brody CD (2021) Collicular circuits for flexible sensorimotor routing. Nature neuroscience 24:1110–1120.

Eckert MA, Keren NI, Aston-Jones G (2010) Looking forward with the locus coeruleius. Science (e-letter).

Edlow BL, Takahashi E, Wu O, Benner T, Dai G, Bu L, Grant PE, Greer DM, Greenberg SM, Kinney HC, Folkerth RD (2012) Neuroanatomic connectivity of the human ascending arousal system critical to consciousness and its disorders. Journal of neuropathology and experimental neurology 71:531–546.

Fusi S, Asaad WF, Miller EK, Wang XJ (2007) A neural circuit model of flexible sensorimotor mapping: learning and forgetting on multiple timescales. Neuron 54:319–333.

Glasser MF, Coalson TS, Robinson EC, Hacker CD, Harwell J, Yacoub E, Ugurbil K, Andersson J, Beckmann CF, Jenkinson M, Smith SM, Van Essen DC (2016) A multi-modal parcellation of human cerebral cortex. Nature 536:171–178.

Glaze CM, Kable J, Gold JI (2015) Normative evidence accumulation in unpredictable environments. eLife 4:e08825.

Glover GH, Li T-Q, Ress D (2000) Image-based method for retrospective correction of physiological motion effects in fMRI: RETROICOR. Magnetic Resonance in Medicine 44:162–167.

Gold JI, Shadlen MN (2007) The neural basis of decision making. Annual review of neuroscience 30:535–574.

Hanks TD, Kopec CD, Brunton BW, Duan CA, Erlich JC, Brody CD (2015) Distinct relationships of parietal and prefrontal cortices to evidence accumulation. Nature 520:220–223.

Harris KD, Thiele A (2011) Cortical state and attention. Nature reviews Neuroscience 12:509–523.

Heinzle J, Wenzel MA, Haynes JD (2012) Visuomotor functional network topology predicts upcoming tasks. The Journal of neuroscience 32:9960–9968.

Hoeks B, Levelt WJ (1993) Pupillary dilation as a measure of attention: A quantitative system analysis. Behavior Research Methods, Instruments & Computers 25:16–26.

Huang Y-Y, Simpson E, Kellendonk C, Kandel ER (2004) Genetic evidence for the bidirectional modulation of synaptic plasticity in the prefrontal cortex by D1 receptors. Proceedings of the National Academy of Sciences 101:3236–3241.

Ito T, Yang GR, Laurent P, Schultz DH, Cole MW (2022) Constructing neural network models from brain data reveals representational transformations linked to adaptive behavior. Nature communications 13:673.

Joshi S, Li Y, Kalwani Rishi M, Gold Joshua I (2016) Relationships between Pupil Diameter and Neuronal Activity in the Locus Coeruleus, Colliculi, and Cingulate Cortex. Neuron 89:221–234.

Kenet T, Bibitchkov D, Dsodyks M, Grinvald A, Arieli A (2003) Spontaneously emerging cortical representations of visual attributes. Nature 425:954–956.

Keren NI, Lozar CT, Harris KC, Morgan PS, Eckert MA (2009) In vivo mapping of the human locus coeruleus. NeuroImage 47:1261–1267.

Kikumoto A, Mayr U (2020) Conjunctive representations that integrate stimuli, responses, and rules are critical for action selection. PNAS 117:10603–10608.

Larsen RS, Waters J (2018) Neuromodulatory Correlates of Pupil Dilation. Frontiers in neural circuits 12:21.

Letzkus JJ, Wolff SB, Luthi A (2015) Disinhibition, a Circuit Mechanism for Associative Learning and Memory. Neuron 88:264–276.

Lin SC, Brown RE, Hussain Shuler MG, Petersen CC, Kepecs A (2015) Optogenetic Dissection of the Basal Forebrain Neuromodulatory Control of Cortical Activation, Plasticity, and Cognition. The Journal of neuroscience 35:13896–13903.

Liu X, de Zwart JA, Scholvinck ML, Chang C, Ye FQ, Leopold DA, Duyn JH (2018) Subcortical evidence for a contribution of arousal to fMRI studies of brain activity. Nature communications 9:395.

Mante V, Sussillo D, Shenoy KV, Newsome WT (2013) Context-dependent computation by recurrent dynamics in prefrontal cortex. Nature 503:78–84.

Maris E, Oostenveld R (2007) Nonparametric statistical testing of EEG-and MEG-data. J Neurosci Methods 164:177–190.

Marzo A, Bai J, Otani S (2009) Neuroplasticity regulation by noradrenaline in mammalian brain. Curr Neuropharmacol 7:286–295.

McGinley Matthew J, David Stephen V, McCormick David A (2015a) Cortical Membrane Potential Signature of Optimal States for Sensory Signal Detection. Neuron 87:179–192.

McGinley MJ, Vinck M, Reimer J, Batista-Brito R, Zagha E, Cadwell CR, Tolias AS, Cardin JA, McCormick DA (2015b) Waking State: Rapid Variations Modulate Neural and Behavioral Responses. Neuron 87:1143–1161.

McGuire JT, Nassar MR, Gold JI, Kable JW (2014) Functionally dissociable influences on learning rate in a dynamic environment. Neuron 84:870–881.

Mesulam MM, van Hoesen GW (1976) Acetylcholinesterase-rich projections from the basal forebrain of the rhesus monkey to neocortex. Brain Research 109:152–157.

Mesulam MM, Changiz G (1988) Nucleus Basalis (CH4) and cortial cholinergic innnervation in the human brain: Observations based on the distribution of acetylcholinesterase and choline acetyltransferase. The Journal of Comparative Neurology 257:216–240.

Mesulam MM, Mufson EJ, Levey AI, Wainer BH (1983) Cholinergic innervation of cortex by the basal forebrain: Cytochemistry and cortical connections of the septal area, diagonal band nuclei, nucleus basalis (substantia Innominata), and hypothalamus in the rhesus monkey. The Journal of Comparative Neurology 214:170–197.

Miller EK, Cohen JD (2001) An integrative theory of prefrontal cortex function. Annual review of neuroscience 24:167–202.

Muller TH, Mars RB, Behrens TE, O’Reilly JX (2019) Control of entropy in neural models of environmental state. eLife 8:e39404.

Munoz DP, Everling S (2004) Look away: the anti-saccade task and the voluntary control of eye movement. Nature reviews Neuroscience 5:218–228.

Murphy PR, O’Connell RG, O’Sullivan M, Robertson IH, Balsters JH (2014) Pupil diameter covaries with BOLD activity in human locus coeruleus. Hum Brain Mapp 35:4140–4154.

Murphy PR, Wilming N, Hernandez-Bocanegra DC, Prat-Ortega G, Donner TH (2021) Adaptive circuit dynamics across human cortex during evidence accumulation in changing environments. Nature neuroscience 24:987–997.

Murty VP, Shermohammed M, Smith DV, Carter RM, Huettel SA, Adcock RA (2014) Resting state networks distinguish human ventral tegmental area from substantia nigra. NeuroImage 100:580–589.

Nadim F, Bucher D (2014) Neuromodulation of neurons and synapses. Current opinion in neurobiology 29:48–56.

Nassar MR, Rumsey KM, Wilson RC, Parikh K, Heasly B, Gold JI (2012) Rational regulation of learning dynamics by pupil-linked arousal systems. Nature neuroscience 15:1040–1046.

O’Reilly JX, Schüffelegen U, Cuell SF, Behrens TE, Mars RB, Rushworth MFS (2012) Dissociable effects of surprise and model updatein parietal and anterior cingulate cortex. PNAS 110:E3660–E3669.

Okazawa G, Kiani R (2023) Neural mechanisms that make perceptual decisions flexible. Annual review of neuroscience 85:191–215.

Pfeffer T, Keitel C, Kluger DS, Keitel A, Russmann A, Thut G, Donner TH, Gross J (2022) Coupling of pupil-and neuronal population dynamics reveals diverse influences of arousal on cortical processing. eLife 11.

Pfeffer T, Ponce-Alvarez A, Tsetsos K, Meindertsma T, Gahnström CJ, van den Brink RL, Nolte G, Engel AK, Deco G, Donner TH (2021) Circuit mechanisms for the chemical modulation of cortex-wide network interactions and behavioral variability. Science Advances 7:eabf5620.

Podvalny E, King LE, He BJ (2021) Spectral signature and behavioral consequence of spontaneous shifts of pupil-linked arousal in human. eLife 10:e68265.

Rasmusson DD (2000) The role of acetylcholine in cortical synaptic plasticity. Behav Brain Res 115:205– 218.

Reimer J, McGinley MJ, Liu Y, Rodenkirch C, Wang Q, McCormick DA, Tolias AS (2016) Pupil fluctuations track rapid changes in adrenergic and cholinergic activity in cortex. Nature communications 7:13289.

Reynolds JNJ, Wickens JR (2002) Dopamine-dependent plasticity of corticostriatal synapses. 15:507–521.

Reynolds JNJ, Hyland BI, Wickens JR (2001) A cellularmechanism of reward-related learning. Nature 413:67–70.

Sara SJ (2009) The locus coeruleus and noradrenergic modulation of cognition. Nature reviews Neuroscience 10:211–223.

Sarafyazd M, Jazayeri M (2019) Hierarchical reasoning by neural circuits in the frontal cortex. Science 364.

Sarter M, Parikh V, Howe M (2009) Phasic acetylcholine release and the volume transmission hypothesis: time to move on. Nature reviews Neuroscience 10:383–390.

Sasaki M, Shibata E, Tohyama K, Takahashi J, Otsuka K, Tsuchiya K, Takahashi S, Ehara S, Terayama Y, Sakai A (2006) Neuromelanin magnetic resonance imaging of locus ceruleus and substantia nigra in Parkinson’s disease. Neuroreport 17:1215–1218.

Schwarz LA, Luo L (2015) Organization of the Locus Coeruleus-Norepinephrine System. Current biology : CB 25:R1051–1056.

Schwarz LA, Miyamichi K, Gao XJ, Beier KT, Weissbourd B, DeLoach KE, Ren J, Ibanes S, Malenka RC, Kremer EJ, Luo L (2015) Viral-genetic tracing of the input-output organization of a central noradrenaline circuit. Nature 524:88–92.

Shadlen MN, Kiani R (2013) Decision making as a window on cognition. Neuron 80:791–806.

Turchi J, Chang C, Ye FQ, Russ BE, Yu DK, Cortes CR, Monosov IE, Duyn JH, Leopold DA (2018) The Basal Forebrain Regulates Global Resting-State fMRI Fluctuations. Neuron.

Turker HB, Riley E, Luh WM, Colcombe SJ, Swallow KM (2021) Estimates of locus coeruleus function with functional magnetic resonance imaging are influenced by localization approaches and the use of multi-echo data. NeuroImage 236:118047.

van den Brink RL, Murphy PR, Nieuwenhuis S (2016) Pupil Diameter Tracks Lapses of Attention. PloS one 11:e0165274.

van den Brink RL, Pfeffer T, Donner TH (2019) Brainstem Modulation of Large-Scale Intrinsic Cortical Activity Correlations. Frontiers in human neuroscience 13.

van den Brink RL, Hagena K, Wilming N, Murphy PR, Buchel C, Donner TH (2023) Flexible sensory-motor mapping rules manifest in correlated variability of stimulus and action codes across the brain. Neuron.

Vetencourt JFM, Sale A, Viegi A, Baroncelli L, De Pasquale R, O’Leary OF, Castrén E, Maffei L (2008) The antidepressant fluoxetine restores plasticity in the adult visual cortex. Science 320:385–388.

Wang L, Mruczek RE, Arcaro MJ, Kastner S (2015) Probabilistic Maps of Visual Topography in Human Cortex. Cerebral cortex 25:3911–3931.

Wang XJ, Yang GR (2018) A disinhibitory circuit motif and flexible information routing in the brain. Current opinion in neurobiology 49:75–83.

Wilming N, Murphy PR, Meyniel F, Donner TH (2020) Large-scale dynamics of perceptual decision information across human cortex. Nature communications 11:5109.

Woolgar A, Jackson J, Duncan J (2016) Coding of Visual, Auditory, Rule, and Response Information in the Brain: 10 Years of Multivoxel Pattern Analysis. Journal of cognitive neuroscience 28:1433–1454.

Yellin D, Berkovich-Ohana A, Malach R (2015) Coupling between pupil fluctuations and resting-state fMRI uncovers a slow build-up of antagonistic responses in the human cortex. NeuroImage 106:414–427.

Zaborszky L, Hoemke L, Mohlberg H, Schleicher A, Amunts K, Zilles K (2008) Stereotaxic probabilistic maps of the magnocellular cell groups in human basal forebrain. NeuroImage 42:1127–1141.

